# Confidence in predicted position error explains saccadic decisions during pursuit

**DOI:** 10.1101/396788

**Authors:** Jonathan D Coutinho, Philippe Lefèvre, Gunnar Blohm

## Abstract

A fundamental problem in motor control is the coordination of complementary movement types to achieve a common goal. As a common example, humans view moving objects through coordinated pursuit and saccadic eye movements. Pursuit is initiated and continuously controlled by retinal image velocity. During pursuit, eye position may lag behind the target. This can be compensated by the discrete execution of a catch-up saccade. The decision to trigger a saccade is influenced by both position and velocity errors and the timing of saccades can be highly variable. The observed distributions of saccade occurrence and trigger time remain poorly understood and this decision process remains imprecisely quantified. Here we propose a predictive, probabilistic model explaining the decision to trigger saccades during pursuit to foveate moving targets. In this model, expected position error and its associated uncertainty are predicted through Bayesian inference across noisy, delayed sensory observations (Kalman filtering). This probabilistic prediction is used to estimate the confidence that a saccade is needed (quantified through log-probability ratio), triggering a saccade upon accumulating to a fixed threshold. The model qualitatively explains behavioural observations on the probability of occurrence and trigger time distributions of saccades during pursuit over a range of target motion trajectories. Furthermore, this model makes novel predictions about the influence of sensory uncertainty and target motion parameters on saccade decisions. We suggest that this predictive, confidence-based decision making strategy represents a fundamental principle for the probabilistic neural control of coordinated movements.

**New & Noteworthy:** This is the first stochastic dynamical systems model of pursuit-saccade coordination accounting for noise and delays in the sensorimotor system. The model uses Bayesian inference to predictively estimate visual motion, triggering saccades when confidence in predicted position error accumulates to a threshold. This model explains saccade probability and trigger time distributions across target trajectories and makes novel predictions about the influence of sensory uncertainty in saccade decisions during pursuit.

## Introduction

The coordination between continuously controlled and discretely triggered movements to achieve a common goal remains a fundamental problem in neuroscience. This coordinated motor control is exemplified in the pursuit and saccadic eye movements that humans perform when tracking moving objects. Pursuit eye movements are continuously controlled to minimize the relative motion of a visual image on the retina (Keller and Heinen, 1991; Lisberger, 2015). Due to noise and delays prevalent in sensorimotor systems (Faisal et al., 2008; van Beers et al., 2002), pursuit trajectory may deviate from and lag behind the true target trajectory (Osborne et al., 2005). Furthermore, pursuit gain in humans is typically less than unity (Diaz et al., 2013a; Ke et al., 2013), resulting in an accumulation of position error during pursuit. As a result, the position of the visual image may drift outside the high acuity foveal region of the retina. Saccades are rapid eye movements that are discretely triggered to reposition the target onto the fovea (Krauzlis, 2004; Orban de Xivry and Lefèvre, 2007). However, the decision process regulating the trigger of saccades during pursuit remains poorly understood. A quantitative model of this decision process has potential to illuminate key principles in the stochastic coordination of continuous and discrete movements for a common goal.

Previous studies have characterized the sensory conditions correlating with saccade trigger during pursuit, though these observations have not yet been synthesized into a mechanistic model. It has been well documented that pursuit can be initiated without saccade occurrence if the target is displaced backwards (step) relative to its motion direction (ramp) such that it crosses the fixation position (target-crossing time) in about 200 ms (Rashbass, 1961). This pattern of step-ramp target motion with target-crossing times slightly earlier or later than this critical 200 ms window tends to evoke saccade with long, variable latencies compared to targets whose motion is in the same direction as the step (Bieg et al., 2015). A similar pattern of saccade probability and trigger time distributions is observed during sustained, steady-state pursuit (de Brouwer et al., 2002b). Thus, the combination of position error and retinal slip are important signals influencing saccades trigger. In one dimensional, horizontal tracking, a documented behavioural correlate of saccade trigger is the negative ratio of position error to retinal slip (i.e. velocity error), which represents the time at which the eye trajectory will cross the target trajectory based on linear extrapolation (de Brouwer et al., 2002b). This ratio, originally named eye-crossing time (T_XE_), will henceforth be referred to as time-to-foveation to parallel the nomenclature of time-to-collision and time-to-contact used in studies of steering and interception (Chang and Jazayeri, 2018; Lee, 1976; Regan and Gray, 2000). Time-to-foveation correlates well with summary statistics of saccade occurrence, but its ability to dynamically predict saccade decisions is limited when retinal slip is close to zero (time-to-foveation goes to infinity) or with two-dimensional target motion (where linear extrapolations of eye and target trajectory commonly fail to intersect). Thus, while existing datasets have outlined the influence of sensory signals on saccade probability and trigger time, a quantitative model explaining trial-by-trial saccade decisions is still missing.

A framework commonly used in modelling decision making under uncertainty is bounded evidence accumulation (Carpenter and Williams, 1995; Gold and Shadlen, 2001; Hanks and Summerfield, 2017). In these models, noisy information is sequentially sampled and the likelihood that this data supports a particular response is integrated over time. Responses are triggered when the accumulated evidence for that response reaches a threshold. Choices, response times, post-decision confidence and neuronal responses in sensorimotor brain areas are consistent with this framework (Fetsch et al., 2018, 2014a, 2014b; Gold and Shadlen, 2007; Schall, 2013, 2000; van den Berg et al., 2016a). Nevertheless, these stochastic decision models have had limited application in oculomotor control, since previous pursuit models relied on deterministic visual motion signals (Krauzlis and Lisberger, 1994a). However, recent Bayesian models successfully simulated the control of pursuit from noisy motion signals (Bogadhi et al., 2013; Orban de Xivry et al., 2013) and predictions about pursuit and saccadic amplitude programming have been validated (Deravet et al., 2018). This provides a novel opportunity to model the stochastic, sensory basis of saccadic decision making during pursuit.

Here we propose a predictive, probabilistic decision mechanism for saccade trigger that explicitly accounts for sensorimotor delays and uncertainties. Retinal position, velocity, and acceleration errors are estimated from noisy, delayed sensory signals through Kalman filtering and predictively extrapolated to overcome sensorimotor delays. Saccades are triggered when saccade confidence accumulates to a threshold. We define saccade confidence as the log probability ratio of the predicted position error being outside the foveal center. This definition has roots in the sequential probability ratio test (Forstmann et al., 2016; Wald and Wolfowitz, 1948) and agrees with the proposition that the term confidence should reflect the probability that a response is appropriate given the observed evidence (Pouget et al., 2016). Furthermore, this framework has a plausible neural basis in probabilistic population coding (Beck et al., 2008; Denève et al., 2007; Ma et al., 2008; Pouget et al., 2003). The model simulates saccade occurrence and trigger time distributions evoked by a wide range of step-ramp target motion trajectories and makes novel predictions about the influence of increased sensory uncertainty (e.g. by blurring the visual target or due to signal-dependent noise). The model illustrates how predictive probabilistic decision making can flexibly coordinate continuously controlled and discretely triggered orienting movements for a common goal. We suggest this stochastic, predictive, confidence-based decision mechanism represents a fundamental principle in the neural control of motor coordination.

## Methods

### Model Overview

The purpose of this model is to explain the sensory basis of saccadic decision making during smooth pursuit eye movements. Using a recent Bayesian model of motion estimation and pursuit dynamics (Orban de Xivry et al., 2013), we develop a novel stochastic evidence accumulation model of saccade triggering. The overall model consists of three interconnected modules: a sensory pathway for predictive state estimation, a decision pathway for evidence accumulation, and a motor pathway implementing the dynamics of eye motion (Fig. 1). The key features of the sensory pathway are recursive Bayesian inference for state estimation (Fig. 1, Kalman Filtering) and prediction through linear motion extrapolation (Fig. 1, Sensory Extrapolation) to compensate for sensorimotor delays and predict future position error. The decision mechanism uses a predictive, probabilistic position error estimate to compute the log-probability ratio that the target to be tracked is left vs right of the fovea (which we define as saccade confidence). A saccade is triggered when leaky accumulation of saccade confidence reaches a threshold value. We use this model to simulate trial-by-trial visual tracking and predict the probability of occurrence and trigger time distributions of saccades evoked by a range of step-ramp target motion trajectories.

**Fig. 1.**
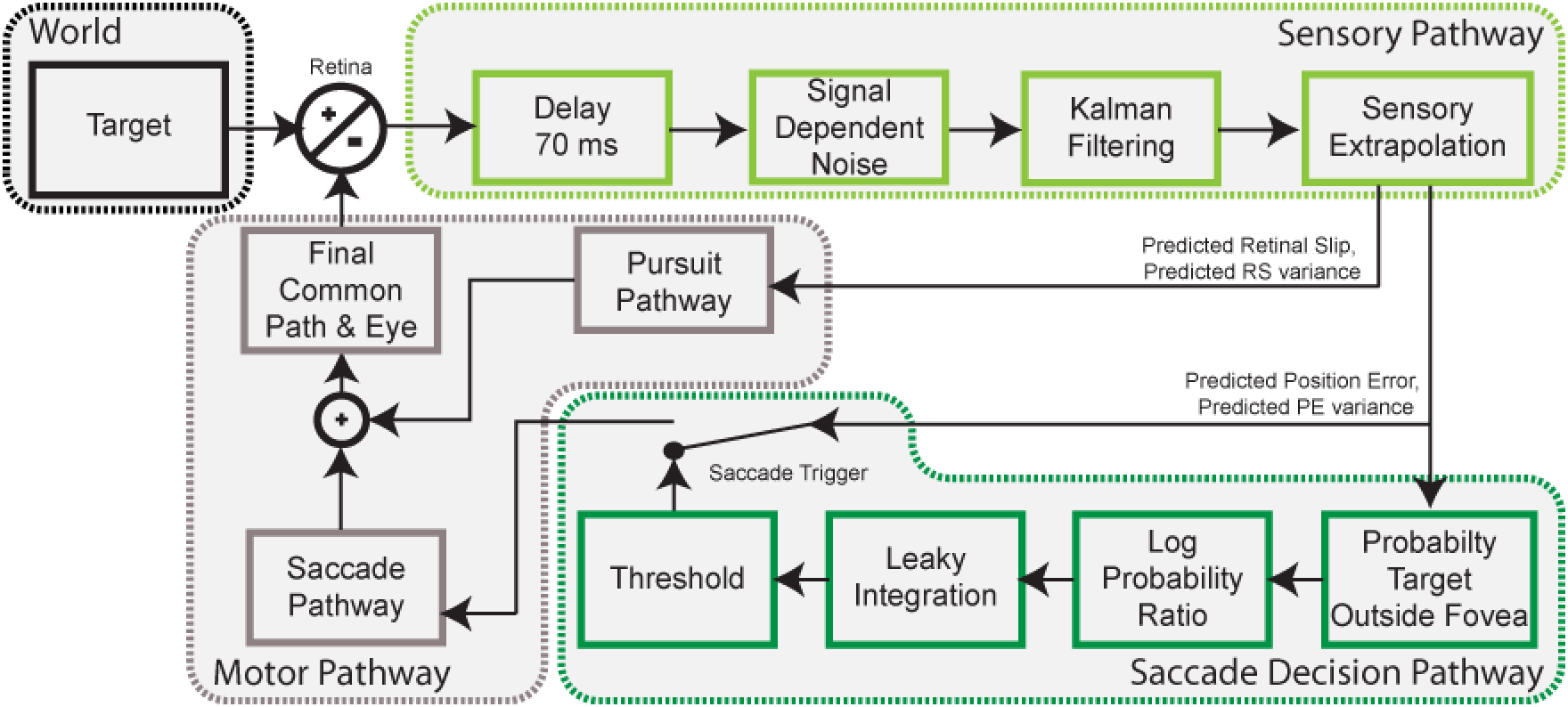
Global overview of model architecture. Sensory pathway (light green): retinal state information is delayed, corrupted by signal-dependent noise, then estimated through Kalman Filtering, and predictively extrapolated to generate predicted position error and predicted retinal slip. Saccade decision pathway (dark green): predicted position error and its associated uncertainty are used to compute the log-probability ratio that the target is left vs right of the fovea. This value, defined as saccade confidence, is accumulated by a leaky integrator to trigger saccades upon threshold crossing. Motor pathway (grey): describes the premotor commands for saccades and pursuit, which are linearly combined in the final common pathway (eye plant) implementing the dynamics of eye motion.

In the detailed model description below, we will denote matrices and vectors in bold, and scalars in unbolded case. Symbols with hat (^) denote estimates of latent variables.

### Sensory Pathway

The sensory pathway is founded on the visual motion processing pathway previously described (Orban de Xivry et al., 2013). The true retinal state, ***δ***^*det*^, is defined as the difference between the position, velocity, and acceleration of the target, ***t***, and the eye, ***e***:

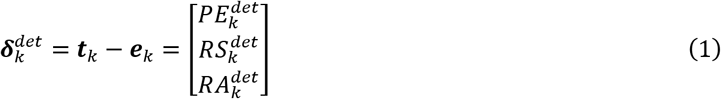

where 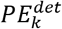 is position error, 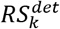 is retinal slip, 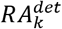 is retinal acceleration, and *k* represents the current discrete time step in the numerical simulation (1 ms).

The observed retinal state, ***δ***^*obs*^, is delayed by 70 ms (Krauzlis and Lisberger, 1994a; Tavassoli and Ringach, 2009) and corrupted by additive and signal-dependent noise:

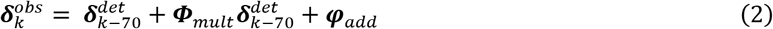

where ***Φ***_*mult*_ and ***φ***_*add*_ are uncorrelated noise covariance matrices with values drawn from 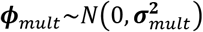 and 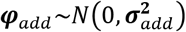 at each time step. The values of noise parameters are listed in Table 1. These terms represent the signal-dependent and baseline noise levels corrupting visual sensory information. Both behavioural and electrophysiological data are consistent with this noise structure in the position and motion information driving saccades and pursuit (Harris and Wolpert, 2006, 1998; Nover et al., 2005; Osborne et al., 2007, 2005; Osborne and Lisberger, 2009; van Beers, 2007).

**Table 1.** Values of noise parameters used in sensory pathway Kalman filtering for retinal position, velocity and acceleration.

| Parameter Name | Parameter Symbol | Parameter Value |
| --- | --- | --- |
| Additive sensory noise variance | $\mathbf{R}^2$ | $\begin{bmatrix} 0.25^2 \\ 7.5^2 \\ 50^2 \end{bmatrix}$ |
| Signal-dependent sensory noise covariance | $\mathbf{D}^2$ | $\begin{bmatrix} 1 & 0 & 0 \\ 0 & 1.5^2 & 0 \\ 0 & 0 & 1 \end{bmatrix}$ |
| Estimated State Variability | $\mathbf{Q}^2$ | $\begin{bmatrix} 0.1 \\ 1 \\ 30 \end{bmatrix}$ |
| Internal Sensory Process Noise | $\boldsymbol{\Omega}^2$ | $\begin{bmatrix} 0.1 \\ 0.3 \\ 10 \end{bmatrix}$ |

Using delayed, noisy observations of image motion, a Bayes-optimal probabilistic estimate of retinal state, 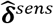, is computed through Kalman filtering (Kalman and Bucy, 1961; Orban de Xivry et al., 2013). This method of recursive Bayesian estimation combines noisy observations (***δ***^*obs*^) with priors of the noise characteristics and dynamics of the world (ie: a generative model) to estimate retinal state. This generative model assumes the evolution of retinal state can be described by an uncorrelated random walk:

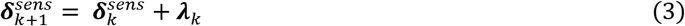

where *λ* is an uncorrelated additive noise with *λ*∼*N*(0, ***Q***^**2**^) representing the state variability, a prior belief about how retinal state changes over time. The generative model also contains priors about the noise characteristics of visual observations of retinal state:

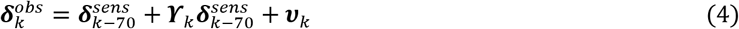

where ***ϒ*** and ***v*** are the expected, uncorrelated signal-dependent and additive noise covariance matrices with ***ϒ***∼*N*(0,***D***^2^) and ***v***∼*N*(0,***R***^2^). These terms represent the brain’s learned estimates of the noise characteristics of sensory observations. We set 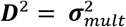 and 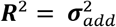. This corresponds to an accurate estimate of noise parameters, although it has been shown an exact knowledge of these values is not crucial (Beck et al., 2012; Orban de Xivry et al., 2013).

Kalman filtering combines the current estimate of retinal state, 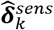, the current noisy observation, 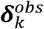, and prior knowledge of noise characteristics, ***R***, ***D***, ***Q***, to optimally estimate retinal state at the next time step (Izawa et al., 2008; Todorov, 2005):

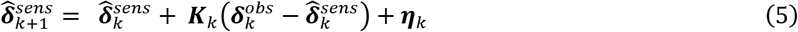

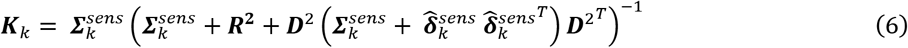

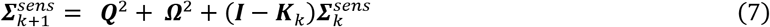

where ***Σ***^*sens*^ is the estimated error covariance of the retinal state estimate, ***K*** is the Kalman gain and ***I*** is the identity matrix. The Kalman gain is calculated and used to weigh incoming sensory evidence according to its relative reliability compared to the current optimal estimate. ***η*** is the internal noise of estimation with ***η***∼*N*(0,*Ω*^2^), which represents variability in the estimation process. The values of Kalman filtering parameters are listed in Table 1.

In the final, predictive stage of the sensory pathway, retinal state is linearly extrapolated through prior knowledge of sensorimotor delays. It has been shown that saccades can accurately foveate moving targets, requiring motion based prediction to extrapolate saccade amplitudes to compensate for the visuomotor delay and ensuing target displacement during saccade execution (Cassanello et al., 2008; de Brouwer et al., 2001; Eggert et al., 2005; Etchells et al., 2010; Guan et al., 2005). Similarly, it has been shown that motion information driving pursuit accounts for both velocity and acceleration, which minimizes instability in pursuit dynamics (Bennett et al., 2007; Krauzlis and Lisberger, 1994b; Lisberger and Westbrook, 1985). Our model hypothesizes that the same predictive position error signal for programming saccade amplitude is used in the mechanism deciding saccade trigger:

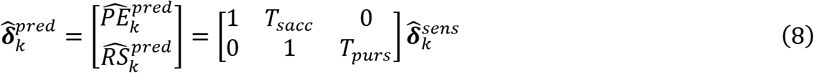

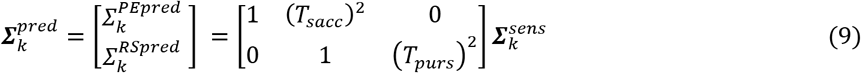

where *T_sacc_* = 125 *ms* and *T_purs_* = 70 *ms* represent the time constant of extrapolation for each respective signal. The rationale for the longer extrapolation time for position error is to account for the additional decision accumulation time, motor delay, and movement duration specific to the saccadic movement system (de Brouwer et al., 2002a).

### Saccade Decision Pathway

The saccade decision pathway is inspired by the sequential probability ratio test and stochastic bounded accumulation models of decision making (Gold and Shadlen, 2007; Wald and Wolfowitz, 1948). This pathway computes the evidence that the target is outside the fovea, i.e. the log-probability ratio of the target being left (*P^Left^*) vs right (*P^Right^*) of the fovea:

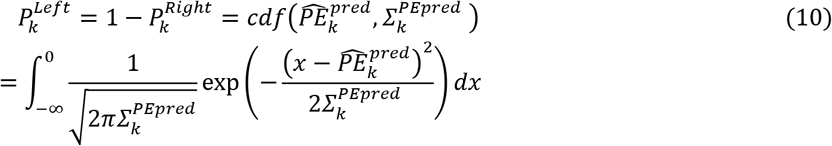

Saccade confidence 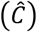 is updated through leaky integration of this evidence:

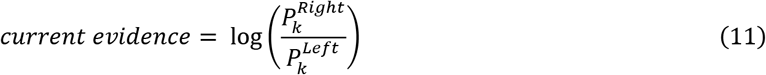

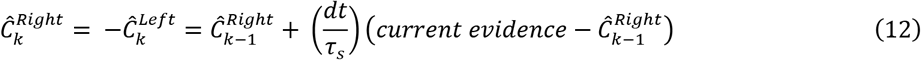

where *τ_s_* = 25 *ms* is the time constant of leaky integration. When predicted PE is close to zero, *P^Right^* and *P^Left^* are close in value, and thus saccade confidence is close to zero. Similarly, as 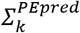 increases with small predicted PE estimates, saccade confidence decreases. With large predicted PE estimates, saccade confidence is less sensitive to 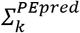. Using leaky integration (Eq. 12), the weight of past evidence exponentially decays at a rate specified by *τ_s_*, biasing decision making towards recent evidence and preventing instantaneous confidence outliers to erroneously trigger a saccade. The estimate of saccade confidence 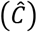 acts as a decision variable triggering saccades when its magnitude exceeds *θ_saccade_* = 4.0. This value corresponds to a 98.2% probability that the target is outside the fovea in a specific horizontal direction. A simplified model of saccade dynamics and refractory period were implemented (see Saccade Motor Pathway below) to illustrate how this decision model can be incorporated into a global framework of oculomotor control.

### Pursuit Motor Pathway

The pursuit motor pathway transforms the predicted retinal slip, 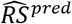, into an oculomotor command that is sent to the premotor system and eye plant. It is adapted from the image velocity motion pathway and positive efferent copy feedback loop used by previous models (Krauzlis and Lisberger, 1994a; Krauzlis and Miles, 1996; Orban de Xivry et al., 2013). The pathway (Fig. 2, Pursuit Motor Pathway, Eq. 13-15) contains a nonlinear transfer function (*G*) to convert the RS input into an oculomotor command, a second-order filter (*H*) to adjust the time course of inputs, and a variable gain element (*A*). The parameters were similar to Krauzlis and Lisberger, 1994; Orban de Xivry et al., 2013.

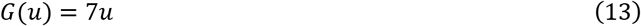

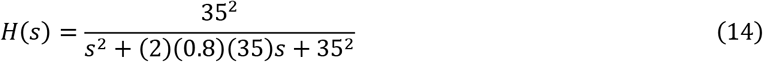

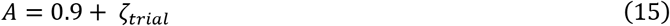

where *s* represents the Laplace variable, and 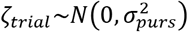, with 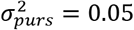. The purpose of this noise term in Eq. 15 is to simulate the trial-by-trial variability in pursuit gain observed in human behavioural responses, thus capturing a realistic level of pursuit variability and consequent trial-by-trial variability in sensory signals based on pursuit performance (Keating and Pierre, 1996; Tabata et al., 2008). While sensory uncertainty is a major source of pursuit variability (Osborne et al., 2005), behavioural evidence suggests further motor related variability influencing pursuit gain that can be dissociated from motion perception (Mukherjee et al., 2015; Rasche and Gegenfurtner, 2009).

**Fig. 2.**
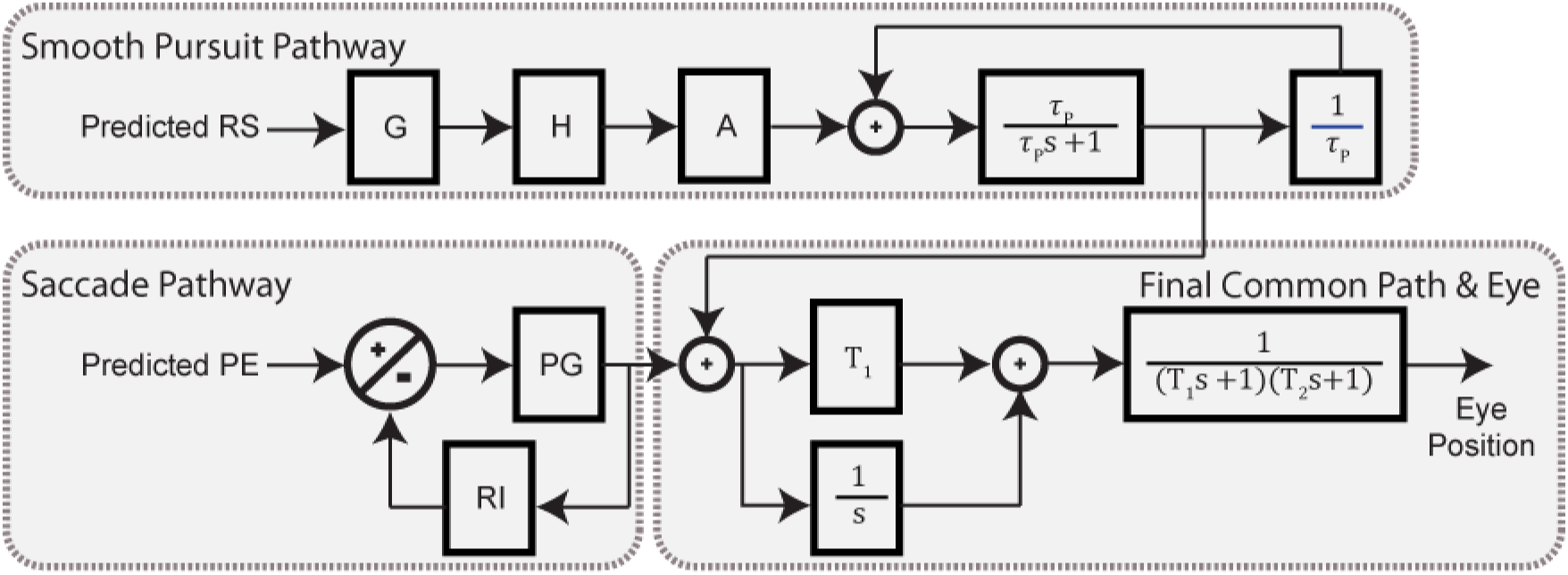
Overview of motor pathway. Pursuit motor commands are continuously generated from predicted retinal slip while saccade motor commands are generated from predicted position error when saccade confidence reaches threshold. Pursuit and saccade motor commands are linearly combined in the final common pathway, representing the neural integration of oculomotor commands for maintaining gaze position and the neuromuscular dynamics of eye motion.

The output of this image velocity motion pathway is sent to a leaky integrator with positive feedback, maintaining eye velocity when 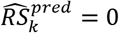. The leaky integrator is characterized by a single time constant, *τ_p_* = 100 *ms*. The positive feedback loop contains a linear gain element, 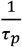, balancing the strength of positive feedback vs leak (Krauzlis and Lisberger, 1994a; Orban de Xivry et al., 2013).

### Saccade Motor Pathway

After saccade confidence reaches the decision threshold, the saccade motor pathway (Fig. 2, Saccade Pathway) transforms a desired saccade amplitude into an oculomotor command that is sent to the premotor system and eye plant. It has been shown that saccade amplitude is programmed by the predicted position error, 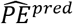, at trigger time (Daye et al., 2014, 2010). The purpose of this saccade model was not to exactly reproduce all aspects of saccade dynamics, but rather to embed a simple, illustrative model of saccade execution within our global framework of oculomotor control. A motor delay of 40 ms was implemented between the time of saccade confidence threshold crossing and the execution of the saccade. The model of saccade dynamics is adapted from the local feedback model and bilateral burst neuron discharge rate pulse generator (Fig. 2, PG) used by (Blohm et al., 2006; Jürgens et al., 1981; Scudder, 1988). In this pathway, the desired saccade amplitude is compared against an internal estimate of executed eye movement through negative feedback from a resettable integrator (Fig. 2, RI), providing an estimate of ongoing motor error without requiring visual feedback. This motor error is sent to a pulse generator, providing a saccadic motor command sent to the premotor system and eye plant. The pulse generator was based on the bilateral burst neuron discharge rate proposed by (Van Gisbergen et al., 1981):

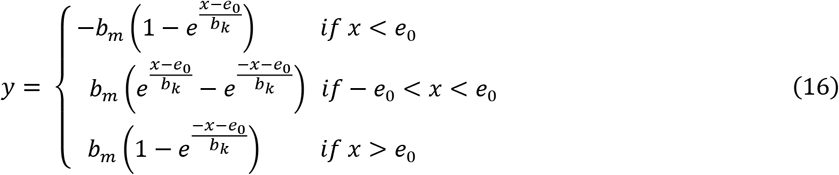

where the input, *x*, is the motor error and the output, *y*, is a saccadic motor command approximating the main sequence relationship between saccade amplitude, peak velocity, and duration (Bahill et al., 1975). The parameters used match Blohm et al., 2006 (ie: *e_0_* = 1 deg; *b_m_* = 600 deg/s; and *b_k_* = 3 deg). Note, that we used an oversimplified saccade generator since saccades were only produced for illustrative purposes and did not affect the smooth pursuit or decision mechanisms. Also, we were only interested in the temporal evolution of trials up until saccade trigger; we thus omitted any variability terms in saccade amplitude which would be required for more realistic catch-up saccade behavior. Furthermore, we avoid speculating on the problem of how retinal signals during the saccade are processed (but see Crevecoeur and Körding, 2017) and how they may influence subsequent pursuit (Goettker et al., 2018; Lisberger, 1998).

### Final Common Motor Pathway and Eye Plant

Oculomotor commands from the pursuit and saccade pathways are linearly added then sent through the premotor system to the eye plant. The premotor system consists of the sum of the motor command (with *gain* = *T*_1_ = 170 *ms*) and its integral, producing the pulse-step innervation pattern required to displace the eye and maintain eccentricity (Robinson, 1973). The eye plant is modelled as an overdamped, second-order system with time constants *T*_1_ = 170 ms and *T*_2_ = 13 ms. No additional motor noise was added to the eye plant, as the moment-by-moment variability in sensory observations and trial-by-trial variability in pursuit gain that we implemented provides a sufficiently realistic distribution of motor variability (Hu et al., 2007; Osborne, 2011; Osborne et al., 2005).

### Simulations

The model was numerically simulated in MATLAB R2018b (Mathworks, Natick, MA, USA) using 1 ms discrete time steps. We simulated eye movements in response to horizontal step-ramp target motions selected to reproduce behavioural investigations of pursuit initiation (Fig. 4,5. Compare to Bieg et al., 2015) and pursuit maintenance (Fig. 6,7,8. Compare to de Brouwer et al., 2002b). For pursuit initiation, we simulated target steps between 1 to 12 degrees in one-degree increments, target velocities of ±10 deg/s and ±20 deg/s (where positive values represent right and negative values represent left), and 100 repetitions at each step-ramp condition. For pursuit maintenance, the initial step-ramp was selected to recross the initial fixation position in 200 ms, thus minimizing initial saccade occurrence. The step sizes used were 2, 4, and 6 degrees with velocity changes of 10, 20, and 30 deg/s respectively in the opposite direction. The second step-ramp was selected using velocity steps of ±10, 20, 40 deg/s with corresponding position steps such to provide a range of target crossing times between −300 ms to 700 ms in 20 ms increments (T_XT_ = −PS/VS), with 50 repetitions per double step-ramp condition. These target motion conditions enable evaluating the influence of T_XE_ on saccade decision making at different target speeds with equal resolution. T_XE_ was calculated at trigger time in saccadic trials, or in smooth trials by taking the time average across the first 400 ms following step-ramp onset.

Simulations of increased sensory noise were implemented using the same double step-ramp conditions while modifying the variance of the additive and multiplicative noise in position estimation to 2^2^ *deg^2^* and 1.5^2^ respectively (Fig. 10). We analyzed the proportion of saccadic trials and saccade trigger time of the first occurring saccade after each step-ramp. Furthermore, we tested the effects of decision parameter variation on these behavioural outcomes (Fig. 11), by simulating visual tracking to a −20 deg/s step-ramp target with varying position steps from −4 to 10 degrees (100 repetitions per step-ramp condition) as either decision accumulator leaky integration time constant was varied between 10 to 75 ms, saccade confidence decision threshold was varied between 3.5 and 4.5, or sensory position extrapolation duration varied from 50 to 250 ms. The code for the full model and all simulations is available on GitHub [https://github.com/Coutinho-J/SaccadeTrigger/].

## Results

This model of oculomotor control relies on Bayesian estimation of stochastic sensory signals and bounded evidence accumulation to trigger saccades upon threshold crossing. The notable novelty of this model is the computation of saccade confidence from a probabilistic prediction of position error. We define saccade confidence as the log-probability ratio that the image is outside the fovea. Saccades are triggered when confidence is leaky accumulated to a threshold value.

### Computation of Saccade Confidence

Saccade confidence is computed from the log-probability ratio of the target being left vs right of the fovea (Eq. 10,11) and leaky integrated over time (Eq. 12). This process is schematically shown in Fig. 3, where a constant predicted PE input is used to illustrate the evolution of saccade confidence over time (although in the normal operation of this model, predicted PE and its uncertainty are time varying based on noisy and evolving retinal state observations). Fig. 3A illustrates the probability density function for a predicted PE of 2 degrees with low uncertainty (purple) and high uncertainty (pink). The probability of the target being right of the fovea, *P^Right^*, is the area under the probability density to the right of zero (denoted in vertical dashed line). The natural logarithm of the ratio between *P^Right^* and *P^Left^* is leaky integrated to compute saccade confidence, whose temporal evolution is illustrated in Fig. 3B. With low magnitude PE, the temporal evolution of saccade confidence is highly sensitive to uncertainty and in the case of an unreliable target even no saccade would be triggered. Fig. 3C illustrates a predicted PE of 4 degrees, with low uncertainty (purple) and high uncertainty (pink). Fig. 3D shows the time evolution of saccade confidence for this PE input, whose time course is much less sensitive to uncertainty because of the larger magnitude PE. Thus, the estimation of saccade confidence accounts for both predicted PE and its uncertainty, but this computation is more sensitive to uncertainty for targets closer to the fovea.

**Fig. 3.**
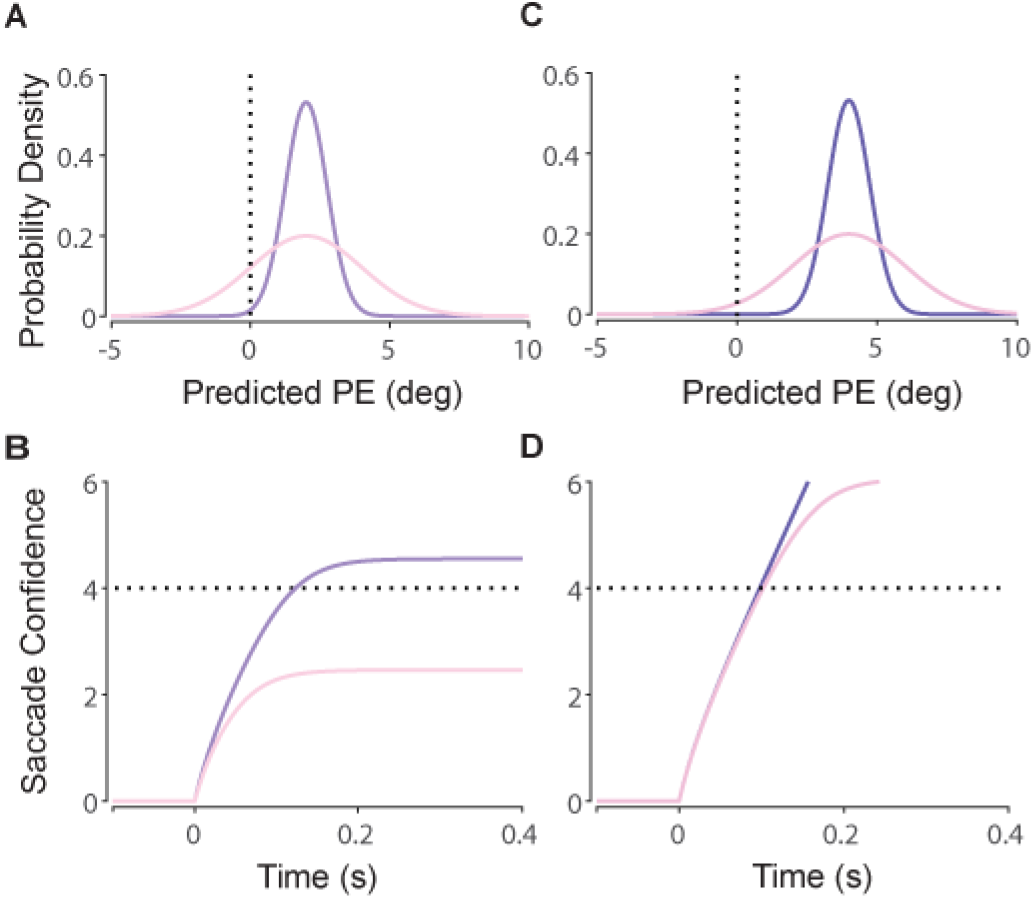
Schematic diagram of the temporal evolution of saccade confidence from constant, probabilistic predicted PE estimates. A) small predicted PE (2 deg) with high (pink) and low (purple) uncertainty. B) This results in small (pink) and large (purple) saccade confidence values. C) Large predicted PE (4 deg) with high (pink) and low (purple) uncertainty. D) This results in little difference in saccade confidence and highly similar decision trigger time.

### Decision Process During Pursuit Initiation

To illustrate how sensory signals influence saccadic decision making, Fig. 4 plots single trial simulations of oculomotor responses during pursuit initiation to two different step-ramp trajectories known to evoke (red trace) and minimize (blue trace) saccade occurrence, as well as a static step of target position (grey trace). The step-ramp trajectories contain an abrupt, simultaneous position displacement and constant velocity shift. This paradigm is commonly used to investigate saccadic decision making during pursuit because it allows the experimenter to precisely control retinal state subsequent to the step-ramp onset (Bieg et al., 2015; de Brouwer et al., 2002b, 2002a; Gellman and Carl, 1991; Rashbass, 1961).

**Fig. 4.**
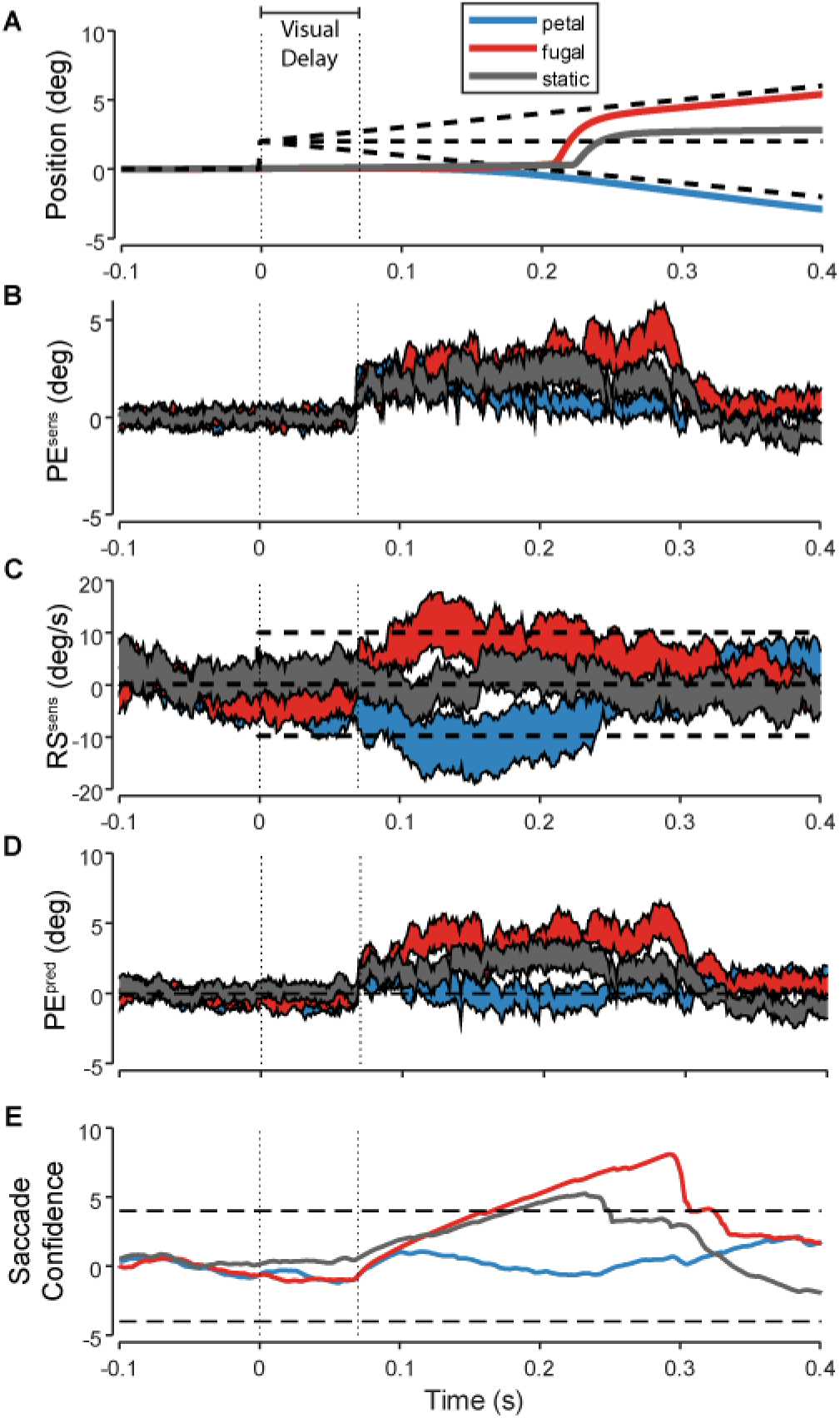
Single trial simulations of pursuit initiation. A) Eye (solid, colored) and target (dashed, black) position over time for a static position step in target (grey) and two step-ramp targets known to evoke (red) and minimize (blue) saccade occurrence. B) Sensory position error over time, estimated from Kalman filtering. Patches represent the estimated value ± its associated uncertainty (std). C) Sensory retinal slip over time, estimated from Kalman filtering. D) Predicted position error over time, estimated by linearly extrapolating sensory position error according to retinal slip. E) Saccade confidence over time, estimated from leaky integrating the log-probability ratio of the target being outside the fovea. Horizontal dashed lines indicate decision threshold.

In our model, retinal state is estimated from noisy and delayed sensory observations through Kalman filtering (Fig. 4B,C) and predictively updated through linear extrapolation (Fig. 4D). This predicted PE is used to compute saccade confidence, which is leaky integrated to trigger saccades upon threshold crossing (Fig. 4E). In Kalman filtering, noisier signals (e.g. RS compared to PE) require more time for accurate estimation, which is comparable to data suggesting a late, asynchronous influence of RS compared to PE in saccade programming (Schreiber et al., 2006). The observation that saccade latency is longer than pursuit latency is explained by the additional temporal accumulator in the saccade decision pathway, intentionally procrastinating decision making to allow for more accurate RS information to accrue and influence the decision. Thus, RS can either enhance or attenuate the evolution of saccade confidence by updating PE away from or towards the fovea, influencing the occurrence and trigger time of saccades.

To reproduce previous findings (Bieg et al., 2015), we simulated oculomotor responses across a range of step-ramp target trajectories and calculated the proportion of saccadic trials and mean saccade trigger time. Fig. 5 plots saccade proportion (Fig. 5A,C) and mean saccade trigger time (Fig. 5B,D) for a variety of step-ramp trajectories where the target velocity either moved towards the fovea (foveopetal) or away from the fovea (foveofugal). This model captures the major trends in behavioural data, replicating the minimization of saccade occurrence when target-crossing time is near 200 ms (Rashbass, 1961), which is the case for a 4 deg step with an target moving at 20 deg/s as in Fig. 5A, or a 2 deg step with the target moving at 10 deg/s as in Fig. 5C. Furthermore, the model replicates the finding of long saccade trigger time for targets with target-crossing times slightly shorter or longer than the 200 ms smooth zone (compare to Fig. 2 in Bieg et al. 2015). Overall, the model reproduces the major trends in saccade trigger during pursuit initiation.

**Fig. 5.**
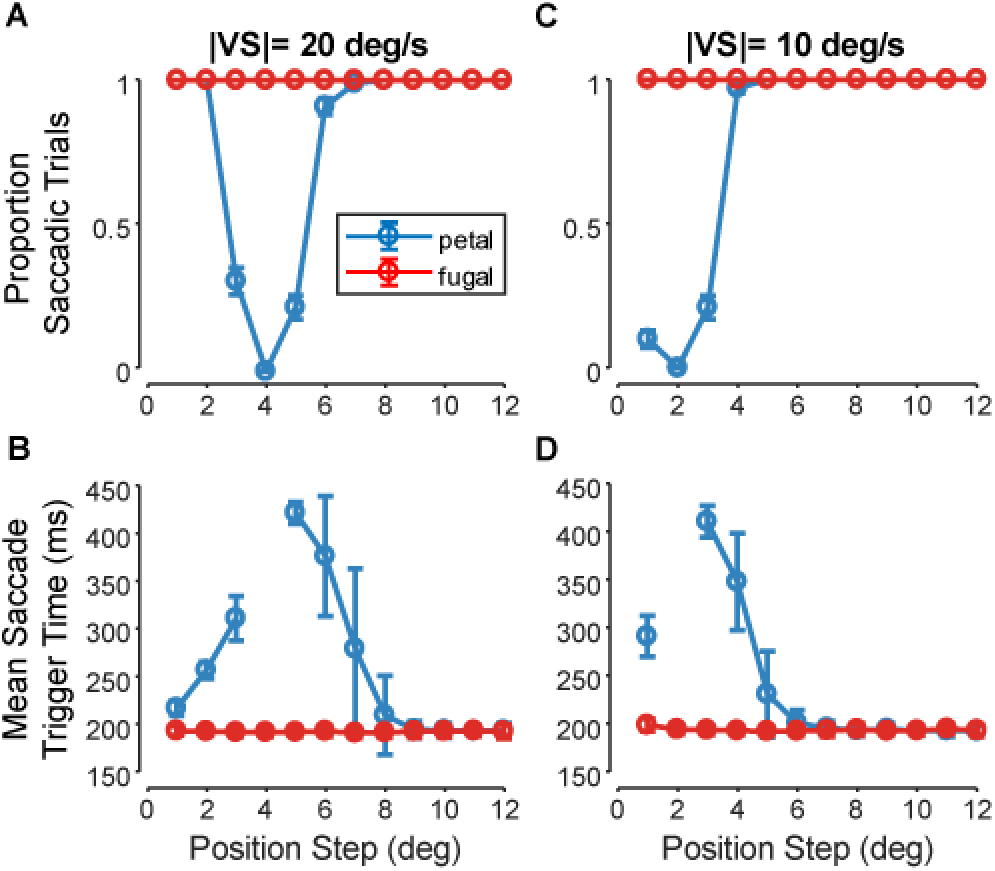
Summary statistics of simulated saccade decisions evoked by step-ramp target motion with position and velocity steps in the same direction (foveofugal, red) or opposite directions (foveopetal, blue). For each step-ramp condition, 100 trial repetitions were simulated. Using a target speed of 20 deg/s and 10 deg/s respectively. A,C) Proportion of trials evoking saccades in the first 450 ms is plotted against position step size. B,D) Mean saccade trigger time ± standard deviation is plotted against position step size.

### Decision Process During Sustained Pursuit

The model was also developed to capture saccade decision making during sustained pursuit using target trajectories with an initial Rashbass paradigm to minimize saccade occurrence followed by a second step-ramp during steady state tracking (de Brouwer et al., 2002b). Fig. 6 illustrates single trial simulations, contrasting smooth and saccadic pursuit in response to two different double step-ramp trajectories. Consistent with saccadic decisions during pursuit initiation, RS estimates that reduce predicted PE result in reduced saccade confidence and subsequent absence of saccade trigger.

**Fig. 6.**
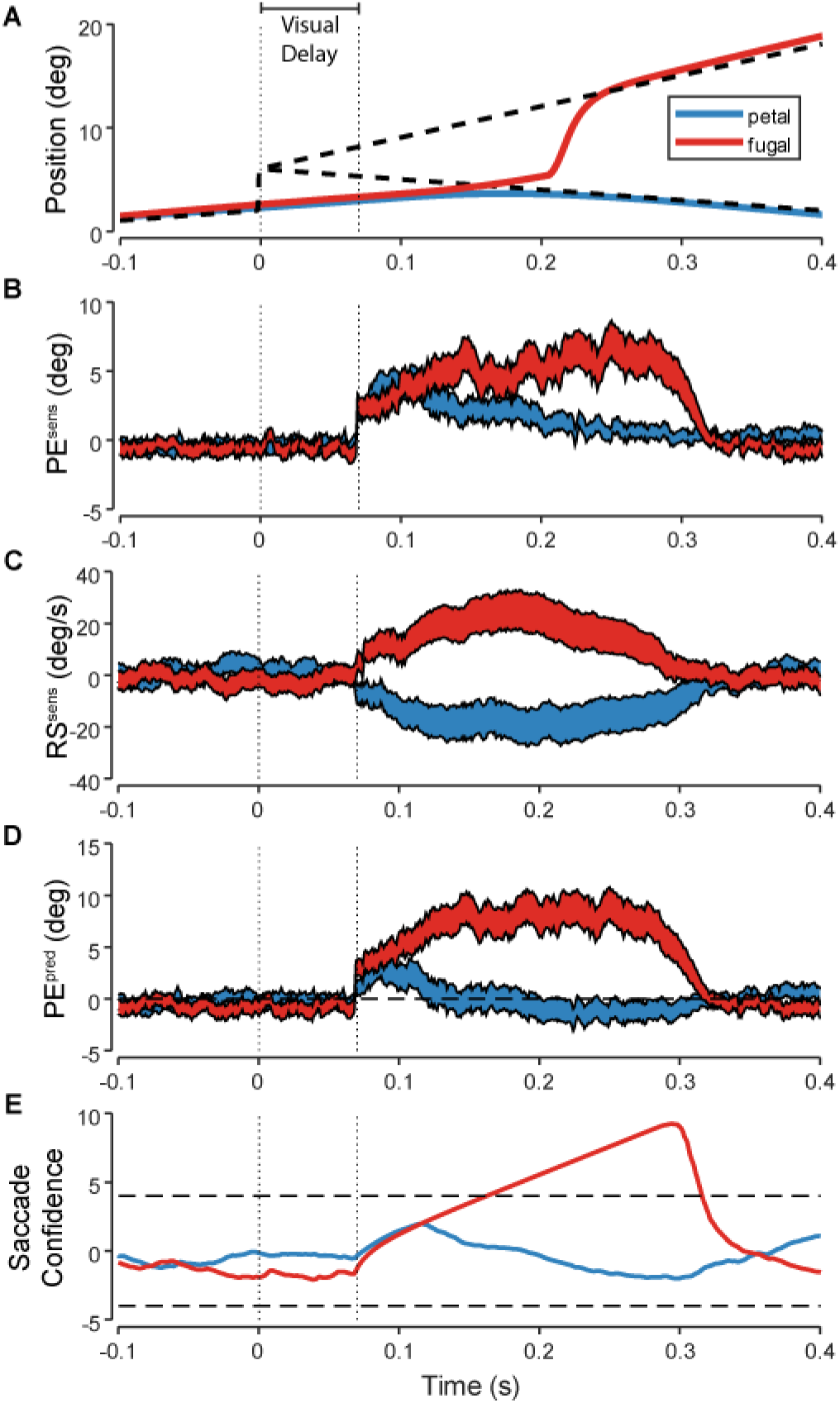
Simulation of single trials of Rashbass step-ramp (not shown) followed by a second step-ramp during steady-state pursuit at time zero. (same conventions as Figure 4).

Across a range of double step-ramp target trajectories, we found that the sensory conditions preceding saccade trigger in simulations were highly comparable to human behavioural data. Fig. 7 illustrates the sensory (Fig.7A) and true (Fig. 7B) values of RS and PE preceding each saccade. The region of this phase plot delimited by solid lines is referred to by de Brouwer and colleagues (2002b) as the smooth zone. The slopes of these lines correspond to single time-to-foveation values. This demonstrates that the model emergently simulates similar correlations between time-to-foveation and human saccade behaviour, namely that saccades are minimized when time-to-foveation is between 40 and 180 ms (the slopes of the lines plotted in Fig. 7). Furthermore, the model simulated saccades with unusually long trigger time occurring when time-to-foveation (T_XE_) was slightly outside this range (Fig. 7, black dots; Fig. 9B). Thus the model also explains saccadic decisions during sustained pursuit.

**Fig. 7.**
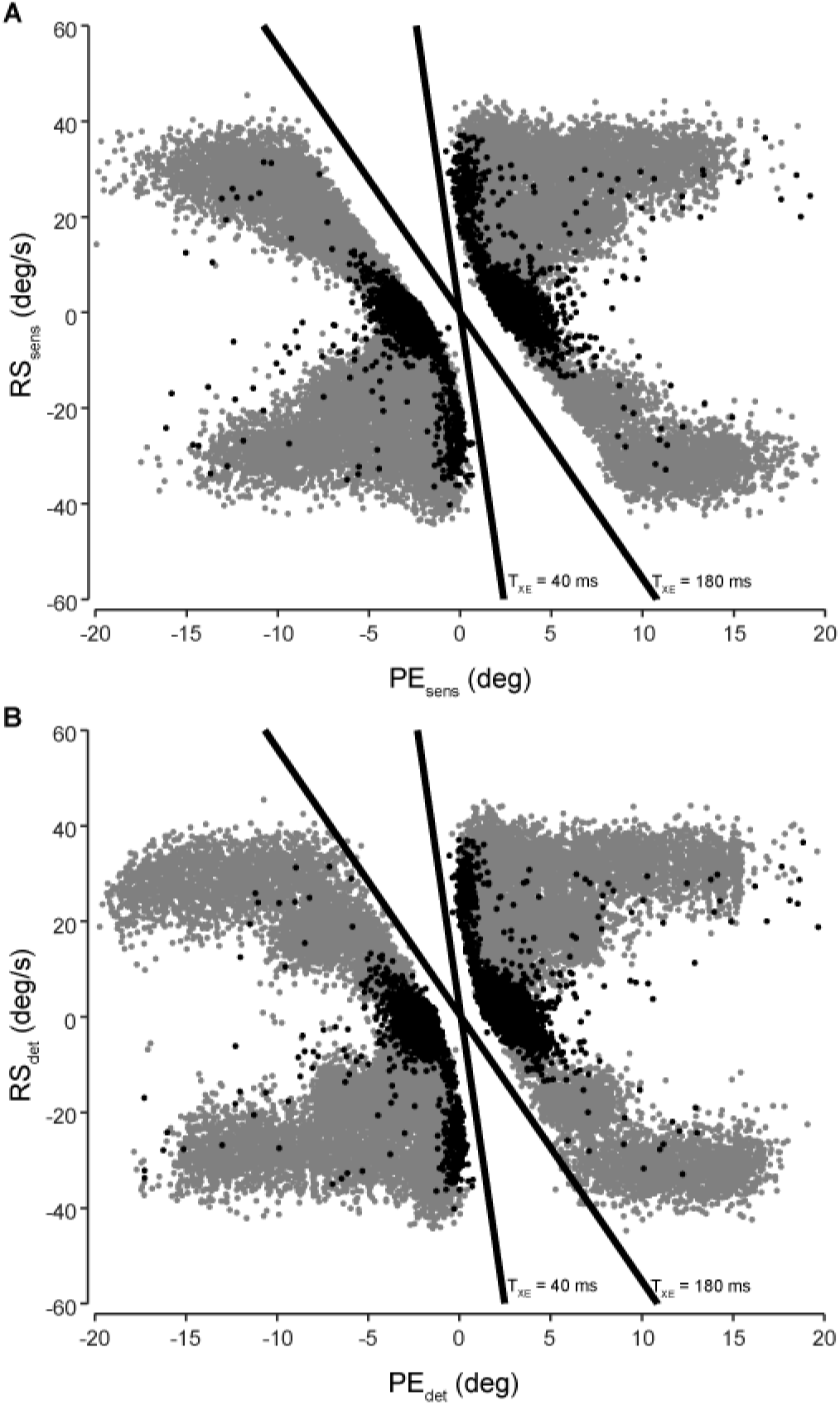
Phase plot illustrating retinal slip vs position error values at time of simulated saccade trigger. A) Sensory RS vs PE. B) True RS vs PE measured 70 ms (visual delay) prior to saccade trigger time. Solid lines represent constant T_XE_ values. Grey dots indicate regular latency saccades, while black dots indicate saccades with trigger time greater than 300 ms. The model emergently simulates the minimization of saccade trigger when T_XE_ is between 40 ms and 180 ms and the occurrence of long latency saccades near this smooth zone.

To further evaluate the relationship between this proposed saccade decision mechanism and previous correlations between saccade trigger and time-to-foveation, we compared the evolution of position error and retinal slip during pursuit with their resulting saccade confidence values. Fig 8 illustrates average saccade confidence contours as a function of position error and retinal slip, along with the evolution of position error and retinal slip during single trials of visual tracking. The onset of step-ramp motion is denoted in circles. During smooth trials (Fig. 8, black trace), position error and retinal slip values lead to subthreshold saccade confidence for the entirety of the trial. During foveofugal target motion (Fig. 8, red trace), position error and retinal slip values lead to high initial saccade confidence and rapid saccade trigger. During some cases of foveopetal target motion (Fig. 8, blue traces), the initial step-ramp results in low saccade confidence which eventually increases to threshold with the subsequent pursuit movement. This explains why long trigger times tend to cluster around the limits of the smooth zone (Fig. 7), since these trials begin with low saccade confidence that ultimately evolves towards threshold confidence much later in the pursuit trajectory.

**Fig. 8.**
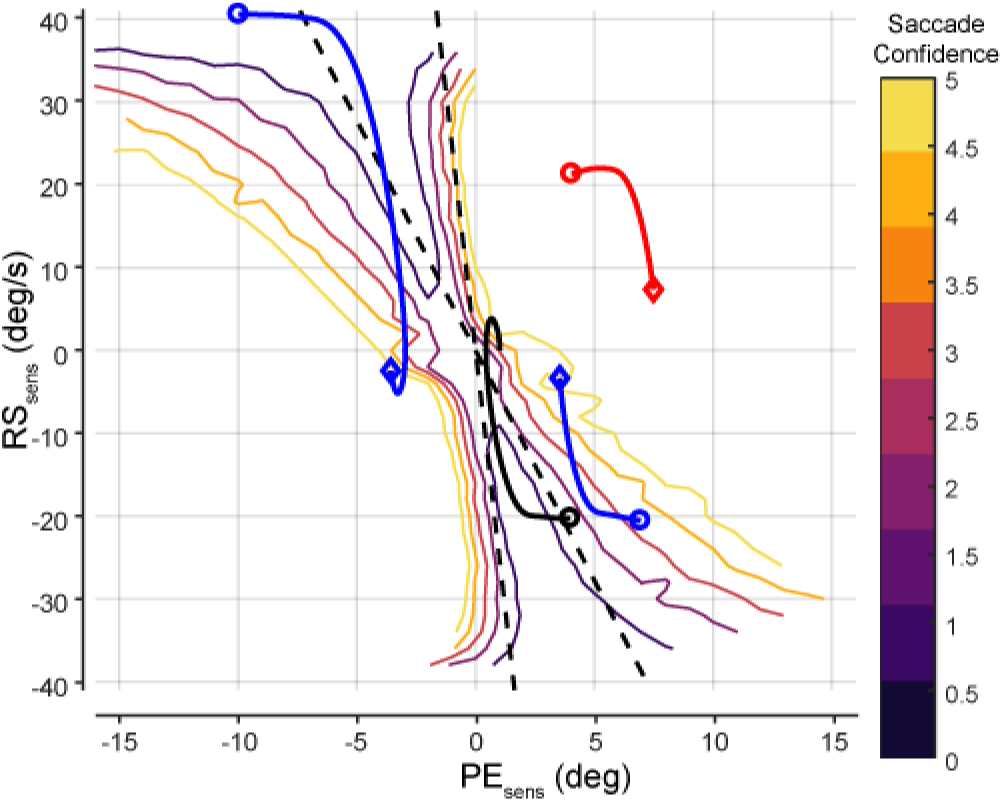
Phase plot illustrating average saccade confidence as a function of position error and retinal slip and the evolution of position error and retinal slip in single trials (onset marked by circle, saccade marked by diamond). Time-to-foveation limits of the smooth zone are plotted as straight lines. Pursuing a foveopetal target with a 2 deg step and −20 deg/s ramp (black) does not trigger a saccade as the combined pursuit and target motion result in low saccade confidence values. Pursuing a foveofugal target with a 2 deg step and 20 deg/s ramp (red) leads to an early saccade since the initial saccade confidence is high and is not reduced by subsequent pursuit. Pursuing a foveopetal target with a 7 deg step and −20 deg/s ramp (blue, bottom right quadrant) results in low initial saccade confidence that eventually increases to threshold during subsequent pursuit, resulting in a long trigger time. Similarly, foveopetal target motion with −10 deg step and 40 deg/s ramp (blue, top left quadrant) results in initial smooth tracking followed by late trigger time after exiting the smooth zone.

### New Model Predictions

Our model makes a series of novel predictions. First, the absolute value of velocity change should correlate with saccade occurrence and trigger time beyond the relationship described by the time-to-foveation parameter. Fig. 9 illustrates saccade proportion (Fig. 9A) and mean saccade trigger time (Fig. 9B) as a function of time-to-foveation (T_XE_), sorted by velocity step for step-ramp motion during sustained pursuit. As the absolute value of the velocity change increases, the range of time-to-foveation values resulting in smooth trials tends to decrease, while saccade trigger time increases near the smooth zone. Therefore, increasing the change in target velocity for a given time-to-foveation value will promote saccade occurrence. However, larger signal-dependent noise due to larger retinal slip results in a slower rise to threshold of saccade confidence and thus longer trigger time near the smooth zone.

**Fig. 9.**
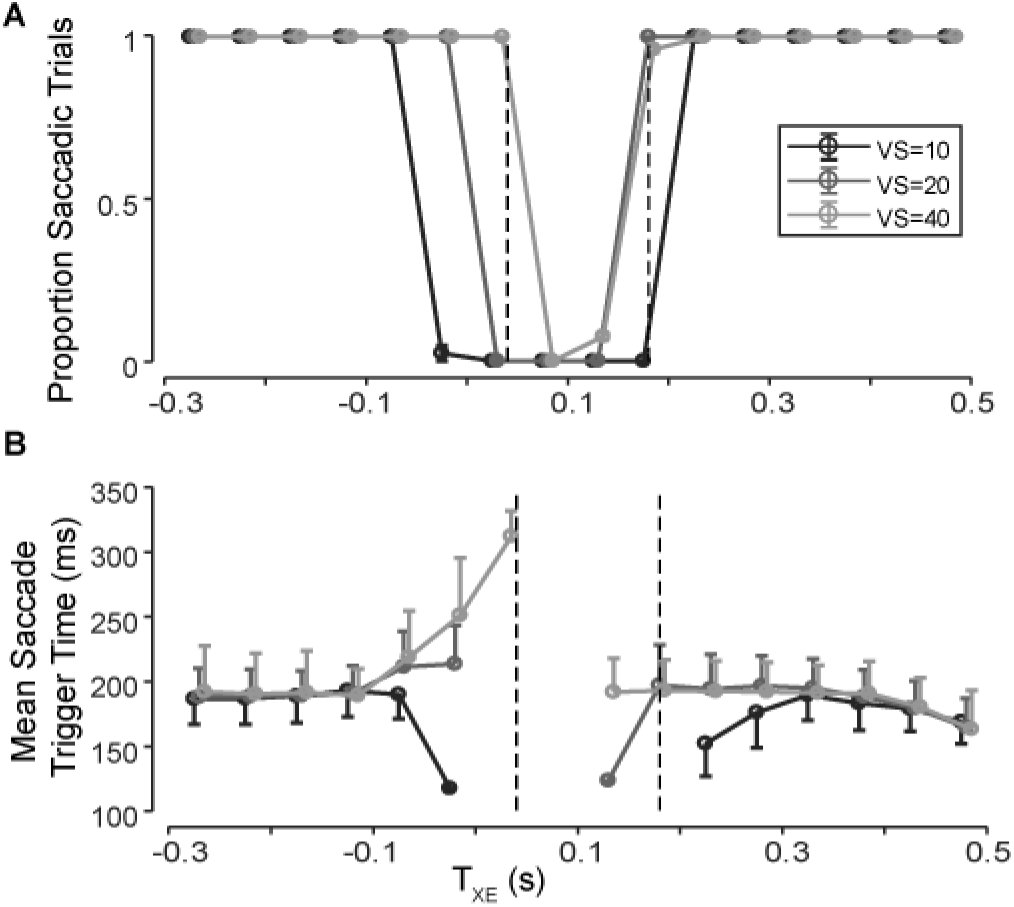
Summary statistics of simulated step-ramp target motion during steady-state pursuit following initial Rashbass step-ramp. A) Proportion of saccadic trials (±sem) plotted against time-to-foveation (T_XE_) and separated by absolute values of change in target speed. For identical time-to-foveation conditions, probability of saccade occurrence is higher for larger changes in target speed exclusively around the smooth zone (defined as time-to-foveation values between 40 and 180 ms, denoted in vertical lines). B) Mean saccade trigger time (±std) plotted against time-to-foveation (T_XE_), separated by absolute value of change in target speed. For identical time-to-foveation conditions, saccade trigger time increases with increasing change in target speed, especially around the smooth zone.

A second novel prediction from this model is that increasing predicted position uncertainty will impact saccade decision making during pursuit. Increasing predicted position noise can be achieved experimentally through Gaussian blurring of the pursuit target and implemented in the model through an increase of the noise corrupting sensory observations. Specifically, we predict a decrease in saccade occurrence and an increase in the variability of saccade trigger time in response to step-ramp target trajectories with time-to-foveation near the smooth zone (Fig. 10 B,E and C,F), but a minimal impact on saccade trigger time distributions for negative time-to-foveation trajectories (corresponding to a position step and velocity change in the same direction) (Fig. 10A,D). This is because the smooth zone corresponds to retinal motion resulting in low predicted PE and saccade confidence is highly sensitive to sensory uncertainty for small predicted PE values (as demonstrated schematically in Fig. 3). Thus, increasing sensory uncertainty should lead to a larger range of tolerable position error estimates that fail to trigger saccades.

**Fig. 10.**
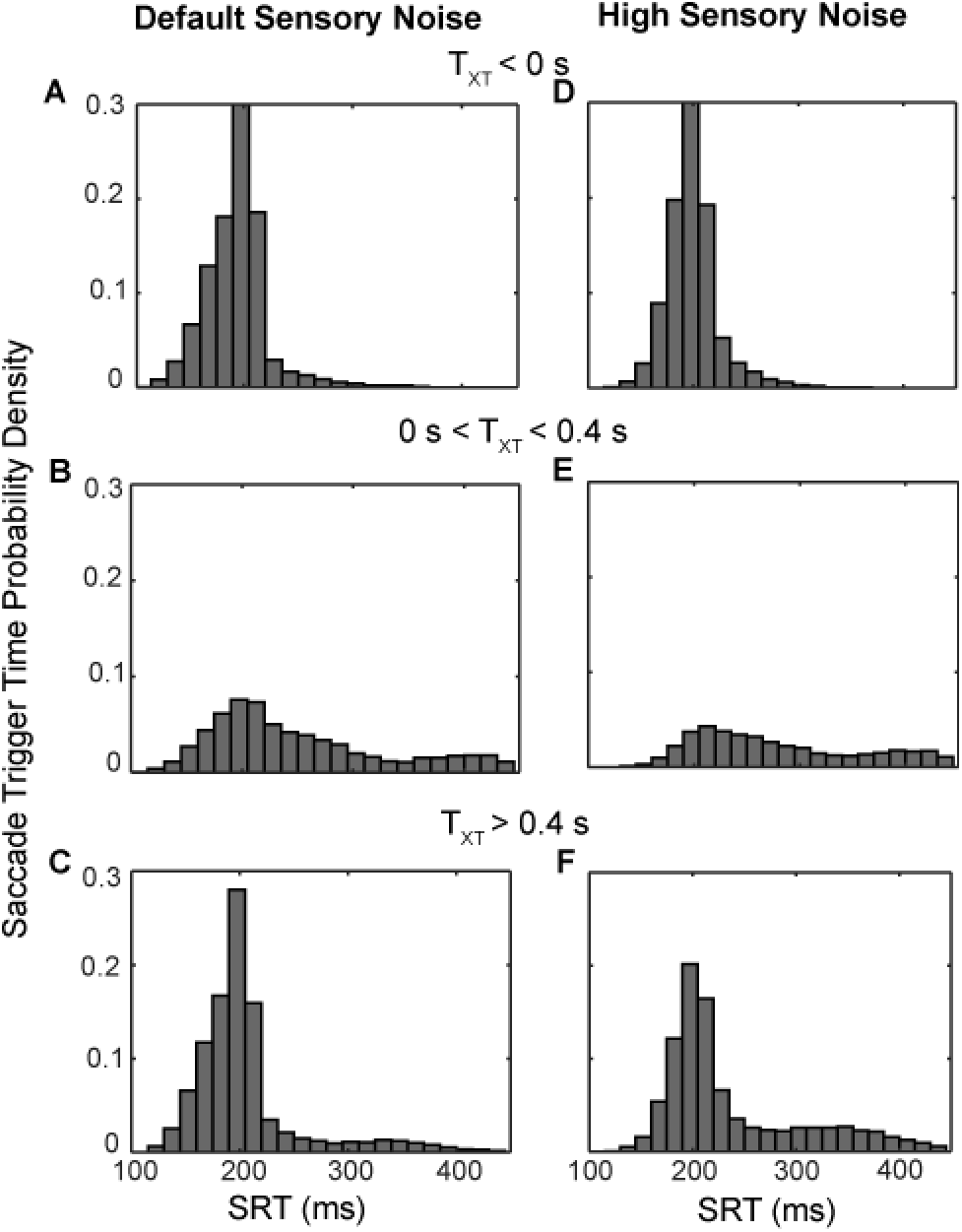
Saccade trigger time probability distributions sorted by target-crossing time for typical (A-C) and unusually high (D-F) sensory position uncertainty grouped by target-crossing time (T_XT_ = −Position Step / Velocity Step). A,D) Increasing sensory noise has minimal influence on saccade trigger time distributions for negative T_XT_ target motion conditions. B,E) This range of T_XT_ values are near the smooth zone, where saccade execution is minimized. With increasing sensory noise there is a further reduction in saccade occurrence and increased variability in saccade trigger time. C,F) With positive T_XT_ values beyond the smooth zone, there is an increase in saccade trigger time variability with increased sensory noise, including an increased probability of late trigger times between 300 ms and 400 ms.

### Effects of Decision Parameter Variations

To illustrate the influence of model parameters within the novel decision mechanism on saccade decision outcomes, we simulated pursuit initiation for step-ramp targets with a velocity of −20 deg/s and varying position steps while varying the value of either the decision accumulator time constant or the saccade confidence decision threshold. With this target speed, saccades are expected to be minimized (i.e. smooth zone) when the target step is 4 degrees (Rashbass, 1961). When increasing the accumulator time constant, instantaneous changes in predicted position error (and its estimated variance) have a weaker influence (i.e. integrated with lower weight) on the evolution of saccade confidence (Eq. 12). This results in a pervasive increase in saccade trigger time regardless of target step size (Fig. 11B) and widening of the smooth zone (Fig. 11A), where larger transient predicted position error values can be tolerated without saccade execution. By increasing the decision threshold, more accumulated evidence of predicted position error is required before triggering saccades. This leads to a slight widening of the smooth zone (Fig. 11C) and slight increase in saccade trigger time around the smooth zone (Fig. 11D). Finally, increasing the motion extrapolation time to compute predicted position error results in a shift and widening of the smooth zone (Fig. 11E). Furthermore, the range of position steps near the smooth zone producing saccades with long trigger time is widened (Fig. 11F), which is caused by the increased uncertainty contributed by the motion pathway (Eq. 9). The same trends hold for saccades during sustained pursuit (not shown).

**Fig. 11.**
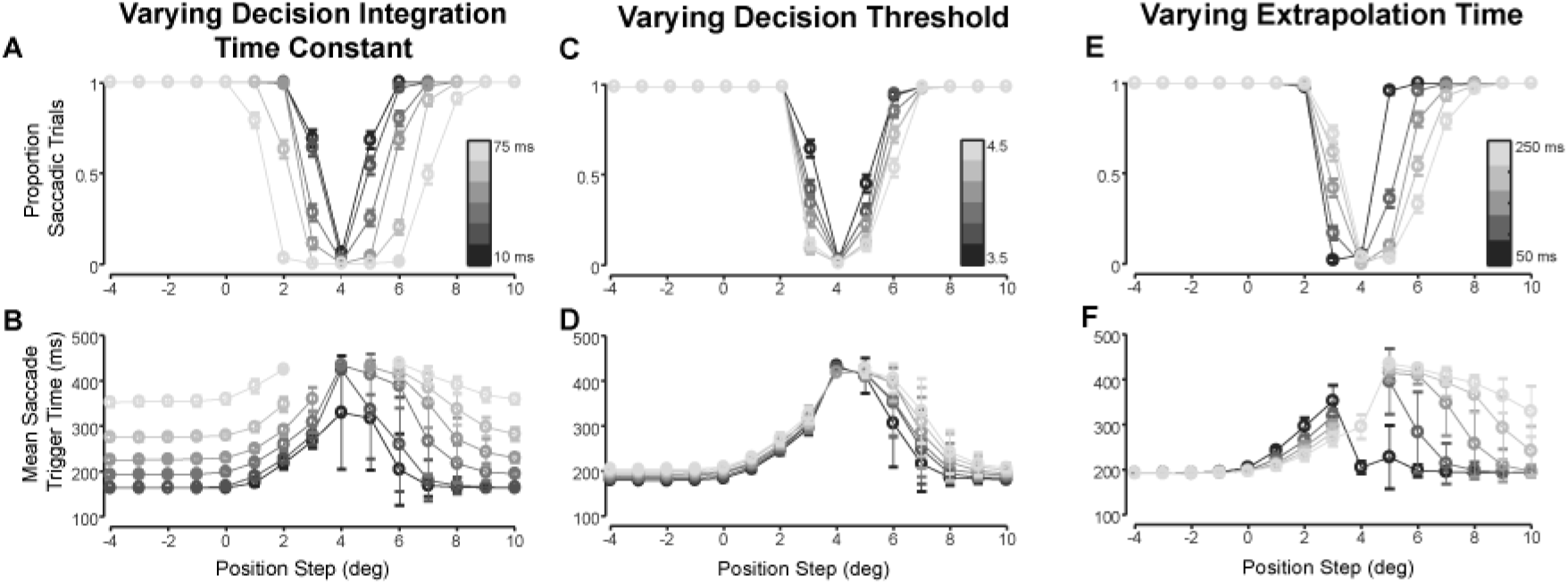
Effects of decision parameter variation on saccade proportion and saccade trigger time during pursuit initiation with a target velocity of −20 deg/s and varying position steps. For these simulations, all other parameters were held at constant, default values. Based on Rashbass (1961), saccade occurrence should be minimized with a position step of 4 deg. A) Increasing the value of the leaky integration time constant in the saccade confidence accumulator (darkest grey corresponds to time constant of 5 ms, lightest grey corresponds to time constant of 50 ms) leads to a widening smooth zone. B) Increasing the value of the leaky integration time constant leads to increased saccade trigger time for all position steps, including far beyond the smooth zone. C) Increasing the value of the saccade confidence decision threshold (darkest grey corresponds to threshold of 3.5, lightest grey corresponds to a threshold of 4.5) results in decreasing saccade proportion near the smooth zone, and a slight widening of smooth zone. D) Increasing the value of the decision threshold leads to slight increases in saccade trigger time only near the smooth zone (darkest grey corresponds to extrapolation time of 50ms, lightest grey corresponds to extrapolation time of 250 ms). E) Increasing sensory extrapolation time shifts the smooth zone. F) Increasing sensory extrapolation time increases saccade trigger time near the smooth zone.

## Discussion

In this model of saccadic decision making during pursuit, confidence in predicted position error triggers saccades upon accumulating to a threshold value. Position error is predictively estimated from noisy and delayed sensory observations to compute the evidence that the target is outside the fovea (saccade confidence, quantified through log-probability ratio). This decision mechanism reproduces the Rashbass paradigm (Rashbass, 1961), as well as the empirical relationships between time-to-foveation and saccade occurrence and trigger time (Bieg et al., 2015; de Brouwer et al., 2002b). The model makes novel predictions about the relationship between target trajectory and saccade behaviour that cannot be accounted by time-to-foveation correlations. Specifically, families of step-ramp target motion with identical time-to-foveation should evoke more long trigger time saccades near the smooth zone (time-to-foveation between 40 and 180 ms, de Brouwer et. al., 2002b) as changes in target speed increases. The model makes further predictions that increasing sensory uncertainty, such as by blurring the visual target (Deravet et al., 2018), will further reduce the probability of saccade occurrence and increase saccade trigger time variability during pursuit, particularly during conditions of low predicted position error (near the smooth zone).

### Model Limitations

This model focuses on the visual basis of saccadic decision making during pursuit without considering the influence of learning and selective attention. In testing saccade behaviour at pursuit initiation, Bieg et al. (2015) randomized the timing, position step size, and motion direction but not motion speed of the target. This could allow for learning speed priors that modify the initial trajectory (and therefore ongoing sensory signals) during the initial pursuit acceleration (Darlington et al., 2017; Dash and Thier, 2013; Ono and Kizuka, 2017). In contrast, de Brouwer et al. (2002b) randomized all aspects of target motion at the second step-ramp, attempting to minimize the influence of learning and expectation, thus providing a more ideal dataset for comparing model performance against. However, selective attention may also present a confounding influence even in the absence of learned expectations. It has been shown that attention is preferentially allocated ahead of pursuit (Chen et al., 2017; Khan et al., 2010) which can preferentially enhance the precision of sensory signals (Poletti et al., 2017). Furthermore, attention may have more direct effects on oculomotor behaviour (Kowler et al., 2014; Zhao et al., 2012), possibly through influencing normative decision thresholds (Lo and Wang, 2006) or accumulation rates (Simen, 2012). Nevertheless, previous modelling studies have demonstrated a dominant influence of sensory variability driving pursuit variability (Osborne et al., 2005), suggesting this model captures the dominant influences on saccade behaviour despite omitting potential cognitive influences. While the mechanisms by which these cognitive factors impact saccadic decision making warrants further investigation, the model successfully illustrates principles of saccade-pursuit coordination and its sensory influences.

Another current limitation that could be addressed in future research is how evolving sensory signals may influence saccade programming after saccade trigger. While it has been clearly shown that saccade amplitude accounts for both position error and retinal slip (de Brouwer et al., 2002a; Eggert et al., 2005; Guan et al., 2005), the influence of retinal slip on saccade amplitude and direction can asynchronously influence the later component of the saccade as demonstrated by curved saccades during two dimensional tracking (Schreiber et al., 2006). Similarly, when saccade targets are abruptly displaced before saccade onset, saccade trajectories may initially aim towards the initial target position then curve midflight towards the final target location (Becker and Jürgens, 1979; Van Gisbergen et al., 1987). This suggests saccade programming is not limited to the information at saccade trigger time, but can still incorporate visual information that only becomes available subsequently (due to delays). Furthermore, during countermanding tasks, erroneously executed saccades have a latency dependent reduction in amplitude (Montagnini and Chelazzi, 2009), suggesting that the stop signal can still influence amplitude programming even after saccade trigger time. These modifications and curvatures in saccade trajectory have been linked to evolving position error estimates after saccade trigger time (McPeek, 2003; Port and Wurtz, 2003; Ramakrishnan et al., 2012). Thus, a potentially interesting extension of this model could investigate how changes in post-trigger predicted position error could influence saccade trajectories.

### Bounded Evidence Accumulation and Confidence Estimation in Decision Making

We have applied the framework of recurrent Bayesian estimation and bounded evidence accumulation towards modelling saccadic decision making in an oculomotor control task. This modelling framework has previously been successfully applied to perceptual decision making, simulating choices, reaction times, and post-decision confidence (Fetsch et al., 2014b). Recently, the concept of confidence estimation has been considering central to the decision process itself (Grimaldi et al., 2015; Kepecs and Mainen, 2012; Meyniel et al., 2015), reflecting the posterior probability that an action is appropriate given the accumulated evidence and potential biases/priors (Pouget et al., 2016). Confidence has been shown to evolve dynamically over the course of evidence accumulation (Dotan et al., 2018), and acts as a critical quantity in multisensory (De Gardelle et al., 2016) and multistage (van den Berg et al., 2016b) decision making. The agreement between model simulations and data suggests the important role of confidence estimation in saccade trigger during pursuit. Thus, we validate previous perceptual decision theory in a novel oculomotor control domain, demonstrating that confidence estimation is also a fundamental principle in motor coordination.

### Generalizability of Saccade Confidence as a Decision Variable

A major feature of the proposed decision mechanism is the generalizability of saccade confidence as a decision variable compared to the previously hypothesized time-to-foveation parameter. Recall that time-to-foveation is defined as the negative ratio between position error and retinal slip. This value rapidly grows to infinity as retinal slip approaches zero. Thus, although the parameter correlates with saccade behaviour after an abrupt change in target motion (de Brouwer et al., 2002b), it is unsuitable as a decision variable for general saccade trigger. In contrast, the mechanism proposed here can explain saccade trigger to both stationary and moving targets.

The model is also compatible with extensions to describe saccade trigger during visual tracking of two-dimensional target motion. This is in contrast to time-to-foveation, which is based on the linear extrapolation and intersection of eye and target trajectories in time; however, eye and target trajectories do not generally intersect in two-dimensional motion. In our framework, a similar saccade confidence estimator may compute the log-probability ratio of the target being above vs below the fovea. These confidence estimators could then be combined to additionally account for vertical and oblique saccade trigger. A challenge would be that independently estimating horizontal and vertical components results in a loss of estimated covariance information. One solution that optimally accounts for covariance is estimating two-dimensional saccade confidence through the Malahanobis distance (De Maesschalck et al., 2000; Mitchell and Krzanowski, 1985) between the fovea and the probabilistic estimate of predicted position error. This measure provides a normalized distance similar to a z-score, uniquely specifying the probability that a reference point (e.g. the fovea) falls within a multivariate distribution (e.g. 2D predicted position error). Thus, the Malahanobis distance could provide an efficient computational mechanism enabling a sequential probability ratio test to estimate saccade confidence in two-dimensional tracking.

### Neurophysiological Implementation

Our model normatively outlines the computational steps underlying saccadic decisions, which has implications on the signals and connectivity of its neural implementation. We thus find it useful to speculate on its potential neurophysiological implementation. Firstly, the model suggests that motion and position information must converge into a predictive position error estimate that ultimately informs the saccade decision process. Motion information is likely supplied by the middle temporal area (MT), since lesions of MT impair the accuracy of saccades to moving targets but not stationary targets (Newsome et al., 1985). Furthermore, electrical microstimulation of MT shifts saccade amplitudes according to the preferred motion direction of the stimulation site and influence saccade trigger time in a target motion dependent manner (Groh et al., 1997). Position information is likely supplied by the superior colliculus (SC), since the spatial locus of activity in the SC corresponds with position error during pursuit (Keller et al., 1996; Krauzlis et al., 2000) and inactivation of the rostral SC (corresponding to foveal position errors) results in stable eye position offsets during pursuit (Hafed et al., 2008). The frontal eye fields (FEF) are characterized as a prediction map for oculomotor control (Crapse and Sommer, 2008) and receive motion information from MT (Ungerleider and Desimone, 1986) and position information from the SC (Lynch et al., 1994). Various neuronal subpopulations of the FEF have been shown to carry the necessary position and velocity signals for predictive saccadic decision making (Bakst et al., 2017; Cassanello et al., 2008). Furthermore, frontal lobe degeneration leads to a dramatic deficit in predictive eye movements (Coppe et al., 2012). Similarly, the lateral intraparietal area (LIP) is involved in the control of both pursuit and saccadic eye movements. LIP has reciprocal connectivity with the FEF and MT, and contains the necessary position and motion information to support predictive saccadic decision making (Berman et al., 1999; Konen and Kastner, 2008; Kurylo and Skavenski, 1991). Additionally, predictive eye position signals have been decoded from the ventral intraparietal area during pursuit (Dowiasch et al., 2016), which may also inform predicted position error. Finally, a cortico-ponto-cerebellar pathway including the dorsolateral pontine nucleus and the cerebellar vermis has sensitivity to both position and velocity information (Dicke et al., 2004) and is involved in controlling both saccades and pursuit (Krauzlis and Miles, 1998; Takagi et al., 2000, 1998). Thus, the computation of predicted position error is likely implemented through this distributed network of cortical and subcortical oculomotor regions.

Secondly, our model suggests neural correlates that integrate probabilistic, predicted PE information over time to compute saccade confidence and evoke saccades. Neuronal activity in LIP during perceptual decision making tasks is consistent with temporal accumulation of evidence supporting the execution of a particular saccade vector (Huk and Shadlen, 2005). In a motion discrimination task, LIP activity is predictive of choice, the relative strength of motion evidence (Shadlen and Newsome, 2001), and post-decision choice confidence (Kiani and Shadlen, 2009). Furthermore, the time that LIP neurons reach a threshold firing rate predicts decision time (Roitman and Shadlen, 2002). Similarly, neuronal activity in the FEF is predictive of saccade choice and timing consistent with evidence accumulation models (Hanes and Schall, 1996). Furthermore, the FEF projects to the caudate nucleus (Cui et al., 2003) and models suggest this corticostriatal pathway terminating in the SC is capable of reading out threshold crossing to trigger saccades (Lo and Wang, 2006). Although further investigation targeting saccade decision signals during pursuit is needed, similar oculomotor networks may be responsible for both prediction and confidence accumulation in this normative decision mechanism.

### Prediction and Confidence Estimation in General Motor Coordination

Coordinated motor decisions are pervasive in our daily lives. For example, eye and hand movements are coordinated when reaching to intercept a moving target. For accurate interception despite sensorimotor delays and movement duration, humans direct their interception movements towards a predicted target location, accounting for the motion of the target and the estimated movement duration (Brenner and Smeets, 2015, 2009; Soechting et al., 2009; Soechting and Flanders, 2008). Furthermore, saccades are often directed to the predicted location of target interception prior to reach initiation (Diaz et al., 2013b; Hayhoe et al., 2003). In a task where monkeys freely choose the timing of manual interception to a moving target, saccades and interceptive reaches are correlated in their initiation timing and endpoint error (Li et al., 2018), suggesting similar decision processes may control both movement types in this coordinated behaviour. Similarly, when humans perform visually guided manual interception, eye movement accuracy strongly predicts the accuracy and timing of interception (Fooken et al., 2016), suggesting both movements are driven by a common internal representation of predicted target trajectory (Goettker et al., 2019). Furthermore, when humans perform interceptive actions, temporal precision is the highest when the interception location was freely chosen compared to pre-specified (Brenner and Smeets, 2015). This is consistent with a decision process that triggers interception when confident in predicted target position. Similarly, during target occlusion, saccades are directed to the predicted location of target reappearance rather than the current predicted location of the occluded target (Orban de Xivry et al., 2009, 2006), further emphasizing that the decision to trigger saccades is based on a confident prediction of target position. Thus, prediction is an important component in ensuring accurate movements, and confidence estimation can evaluate the appropriateness of particular actions given the uncertainty of predictions.

Similar coordinated movements occur in a variety of animal species, who may also utilize similar decision principles in coordinating movements from delayed and uncertain sensory signals. For example, flies (*Drosophila*) in visually guided flight perform coordinated smooth and saccadic turning in order to accurately orient towards their flight goal (Mongeau and Frye, 2017). Models including temporal integration of motion information have been employed to explain this rapid decision making (Mongeau et al., 2019, 2018), but it has not yet been investigated how uncertainty in visual motion may influence these decisions. As another example, zebrafishes integrate specific visual features in the perceptual decision of prey recognition, which can initiate hunting routines such as convergent saccades, orienting turns, and capture swims (Bianco et al., 2011). The visual features important to prey detection have been described (Semmelhack et al., 2014) and the neural circuits mediating prey recognition and the initiation of hunting have been localized to non-overlapping populations in the optic tectum (Bianco and Engert, 2015; Dunn et al., 2016; Fajardo et al., 2013). However, it remains unknown how sensory uncertainty is handled within these decision circuits when initiating discretely triggered components of the hunting routine. These species are noteworthy for the wealth of experimental techniques available for investigating detailed neural circuits underlying sensorimotor behaviour (Gahtan and Baier, 2004; Olsen and Wilson, 2008; Orger, 2016; Stowers et al., 2017). Thus, principles from our model could be used to guide research questions with these animal models (Clemens and Murthy, 2017) to investigate cellular and circuit mechanisms of prediction and confidence estimation for general motor coordination.

## Conclusion

Our model of saccade decision making during pursuit handles the constraints of delay and signal-dependent noise in the oculomotor system while reproducing established trends in saccade occurrence and trigger time across a range of target motion trajectories. The model illustrates how discretely triggered orienting movements like saccades can be coordinated during continuously controlled pursuit through predictive, probabilistic evidence (confidence) accumulation. We suggest that this framework of prediction and confidence estimation represents a fundamental principle in stochastic decision making for sensorimotor coordination.

## Acknowledgements

This work was supported by the National Sciences and Engineering Research Council (NSERC, Canada) and the Canada Foundation for Innovation (CFI).

